# Meta-analysis of RNA-seq studies reveals genes responsible for life stage-dominant functions in *Schistosoma mansoni*

**DOI:** 10.1101/308189

**Authors:** Zhigang Lu, Matthew Berriman

## Abstract

**Background:** Since the genome of the parasitic flatworm *Schistosoma mansoni* was sequenced in 2009, various RNA-seq studies have been conducted to investigate differential gene expression between certain life stages. Based on these studies, the overview of gene expression in all life stages can improve our understanding of *S. mansoni* genome biology.

**Methods:** publicly available RNA-seq data covering all life stages and gonads were mapped to the latest *S. mansoni* genome. Read counts were normalised across all samples and differential expression analysis was preformed using the generalized linear model (GLM) approach.

**Results:** we revealed for the first time the dissimilarities among all life stages. Genes that are abundantly-expressed in all life stages, as well as those preferentially-expressed in certain stage(s), were determined. The latter reveals genes responsible for stage-dominant functions of the parasite, which can be a guidance for the investigation and annotation of gene functions. In addition, distinct differential expression patterns were observed between adjacent life stages, which not only correlate well with original individual studies, but also provide additional information on changes in gene expression during parasite transitions. Furthermore, thirteen novel housekeeping genes across all life stages were identified, which is valuable for quantitative studies (e.g., qPCR).

**Conclusions:** the metaanalysis provides valuable information on the expression and potential functions of *S. mansoni* genes across all life stages, and can facilitate basic as well as applied research for the community.

## Introduction

Schistosomes are parasite flatworms that infect more than 240 million people worldwide, and cause up to 200,000 deaths every year (GBD 2015 Mortality and Causes of Death Collaborators 2016). There is currently no vaccine, and only one putatively-used drug. To accelerate the elimination of this parasite, it is important to understand its biology, in particular its genome biology. First sequencing of schistosome genomes were accomplished in 2009, for two of the most common *Schistosoma* species, *S. mansoni* (Berriman et al. 2009) and *S. japonicum* (The Schistosoma japonicum Genome Sequencing and Functional Analysis Consortium 2009). Afterwards, an improved version of *S. mansoni* genome was achieved (Protasio et al. 2012). The genome information provides a basis for many high throughput approaches, among which RNA sequencing (RNA-seq) has been developed and exploited for determining gene expression in recent years.

Due to the complexity of schistosome life cycle, as well as the high sequencing cost in the past, previous RNA-seq studies were mainly focused on certain life stages, such as the larvae (Wang et al. 2013; Protasio et al. 2012), the adult stage (Lu et al. 2016; Anderson et al. 2015), or based on certain experimental conditions, e.g., UV-radiation treatment (Collins III et al. 2013). While we can get valuable information from the partial comparisons, an overall estimation of gene expression during schistosome development is missing. On the other hand, expression profiling in all life stages is also able to improve functional annotation of the genome, especially for stage-specifically expressed genes and hypothetical genes (Elias et al. 2009).

To obtain the information on gene expression changes in different life stages, we performed a comprehensive meta-analysis on published RNA-seq studies in *Schistosoma mansoni*. We identified genes that might account for the dominant function of the parasite in specific life stage(s), which will be important for understanding the biology of schistosomes and the process of parasitism. Further explored differential gene expression during life stage transitions, as well as those ubiquitously- and/or abundantly-expressed genes can benefit basic and applied research for the community.

## Methods

### Implemented life stages and RNA-seq sequence data

All sequence data were obtained from ENA (http://www.ebi.ac.uk/ena), originating from four published studies. The accession numbers and sample types were summarised in Table 1.

**Table 1.**
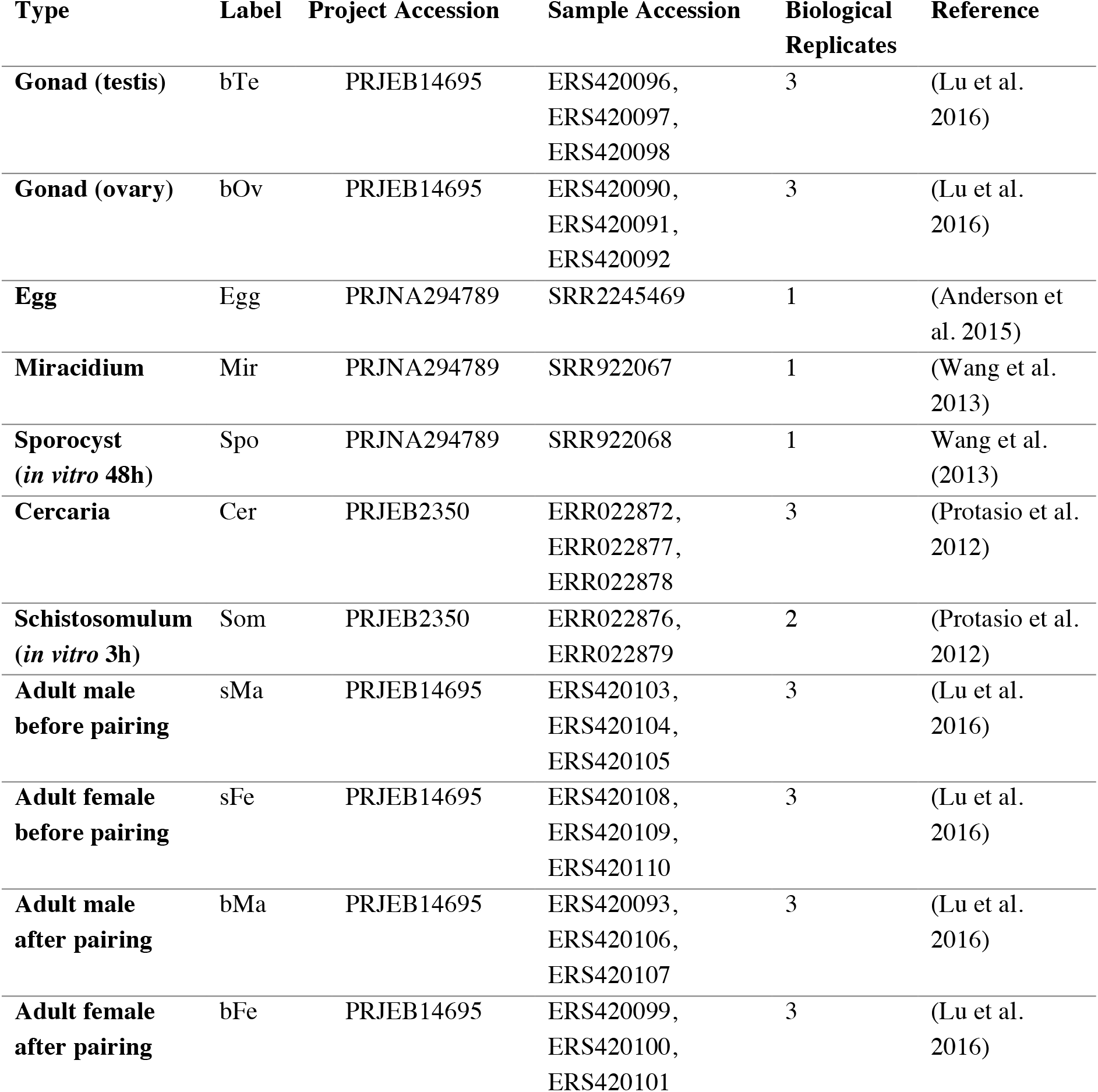
Summary of datasets and samples analysed in this study

| Type | Label | Project Accession | Sample Accession | Biological Replicates | Reference |
| --- | --- | --- | --- | --- | --- |
| Gonad (testis) | bTe | PRJEB14695 | ERS420096,<br>ERS420097,<br>ERS420098 | 3 | (Lu et al. 2016) |
| Gonad (ovary) | bOv | PRJEB14695 | ERS420090,<br>ERS420091,<br>ERS420092 | 3 | (Lu et al. 2016) |
| Egg | Egg | PRJNA294789 | SRR2245469 | 1 | (Anderson et al. 2015) |
| Miracidium | Mir | PRJNA294789 | SRR922067 | 1 | (Wang et al. 2013) |
| Sporocyst (in vitro 48h) | Spo | PRJNA294789 | SRR922068 | 1 | Wang et al. (2013) |
| Cercaria | Cer | PRJEB2350 | ERR022872,<br>ERR022877,<br>ERR022878 | 3 | (Protasio et al. 2012) |
| Schistosomulum (in vitro 3h) | Som | PRJEB2350 | ERR022876,<br>ERR022879 | 2 | (Protasio et al. 2012) |
| Adult male before pairing | sMa | PRJEB14695 | ERS420103,<br>ERS420104,<br>ERS420105 | 3 | (Lu et al. 2016) |
| Adult female before pairing | sFe | PRJEB14695 | ERS420108,<br>ERS420109,<br>ERS420110 | 3 | (Lu et al. 2016) |
| Adult male after pairing | bMa | PRJEB14695 | ERS420093,<br>ERS420106,<br>ERS420107 | 3 | (Lu et al. 2016) |
| Adult female after pairing | bFe | PRJEB14695 | ERS420099,<br>ERS420100,<br>ERS420101 | 3 | (Lu et al. 2016) |

### Sequence mapping and reads counting

Sequences were mapped to *S. mansoni* genome V5.2 using STAR (Dobin et al. 2013) (v2.4.2a) with the parameter *alignIntronMin* set to 10 for all samples except egg, or using HISAT2 (Kim et al. 2015) (v2.1.0) for the egg sample. Counts per gene were summarised with featureCounts (Liao et al. 2014) (v1.4.5-p1) on the latest annotation (GeneDB (www.genedb.org; data accessed 10/07/2017) and used for downstream analysis.

### Principle Component Analysis (PCA) and Hierarchical Clustering

Sample dissimilarities were revealed by PCA using the R (https://www.r-project.org) (v3.3.2) package DESeq2 (Love et al. 2014) (v1.14.1), with the parameter *ntop* set to the total number of genes. Plotting of PCA data were performed using default settings. Sample distance matrix was calculated using the sample package. Hierarchical clustering of all genes was performed using the *hclust* function and the “ward.D2” method. Heat maps were generated using the gplots (https://cran.r-project.org/web/packages/gplots/index.html) (v2.2.1) package.

### Normalisation and differential expression analysis

Read counts were imported into edgeR (Robinson et al. 2010) (v3.16.5) and normalised across all samples by the Trimmed Mean of M-values (TMM) method (Robinson et al. 2010) using the function *calcNormFactors()*. Differential gene expression were analysed with the Generalized Linear Models (GLM) approach using the functions *glmFit()* and *glmLRT()*.

### Determination of most abundantly-expressed, and stage(s)-preferentially expressed genes

To obtain most abundantly-expressed genes in all life stages, mean expression values (before normalisation) of genes in all samples excluding gonads were ranked and the top twenty were listed. Life stage(s)-preferentially expressed genes were defined as significant higher expression in one sample/group than in the rest of samples. This was calculated by using the GLM approach and setting False Discovery Rate (FDR) < 0.01. Further manual filtration was performed to select genes with higher expression in certain stage(s) than in any other stage.

### Gene Ontology (GO) terms enrichment and KEGG pathway mapping

GO terms enrichment analysis was performed with PANTHER (Mi et al. 2013) (v12.0; analysed on 24/08/2017) using Bonferroni correction and P < 0.05 as threshold. Significantly enriched Biological Processes were plotted according to the P-values. For pathway mapping, protein sequences of all genes were mapped to the KEGG pathway database (Kanehisa and Goto 2000) using KAAS (http://www.genome.jp/kaas-bin/kaas_main; analysed on 08/08/2017; program: GHOSTZ, alignment method: BBH).

### Identification of housekeeping genes

Housekeeping genes were identified with the GLM approach which compared all life stages (excluding gonads) to bFe, as an arbitrarily selected reference. Candidate housekeepers were selected from those with no significant difference (FDR > 0.05) and in addition fold-difference < 1.5. Further testing of their suitability was performed by calculating the stability value among all samples using the R package Normfinder (Andersen et al. 2004) (v05/01-2015).

### Correlations with original studies

To check the correlations between meta-analysis and original studies, differentially expressed genes (FDR cut-off 0.01) were selected from a specific comparison and corresponding log2-based fold-change (log_2_FC) values from both analyses were used to calculated Pearson correlation coefficient.

## Results

### Dissimilarities between all life stages and stage-associated expression profiling

Across all life-cycle stages and all experiments, the proportion of genes showing convincing expression (RPKM > 1) varied from 58% to 83% (Table 2), with eggs displaying the smallest repertoire of expressed genes. By Principle Component Analysis (PCA) (Fig. 1A) several clusters were identified: bOv, bTe, Egg, Mir-Spo, Cer-Som, sFe-sMa-bMa, and bFe. This set of dissimilarities were confirmed by sample distance matrix (Fig. 1B) and hierarchical clustering analysis (Fig. 1C).

**Figure 1.**
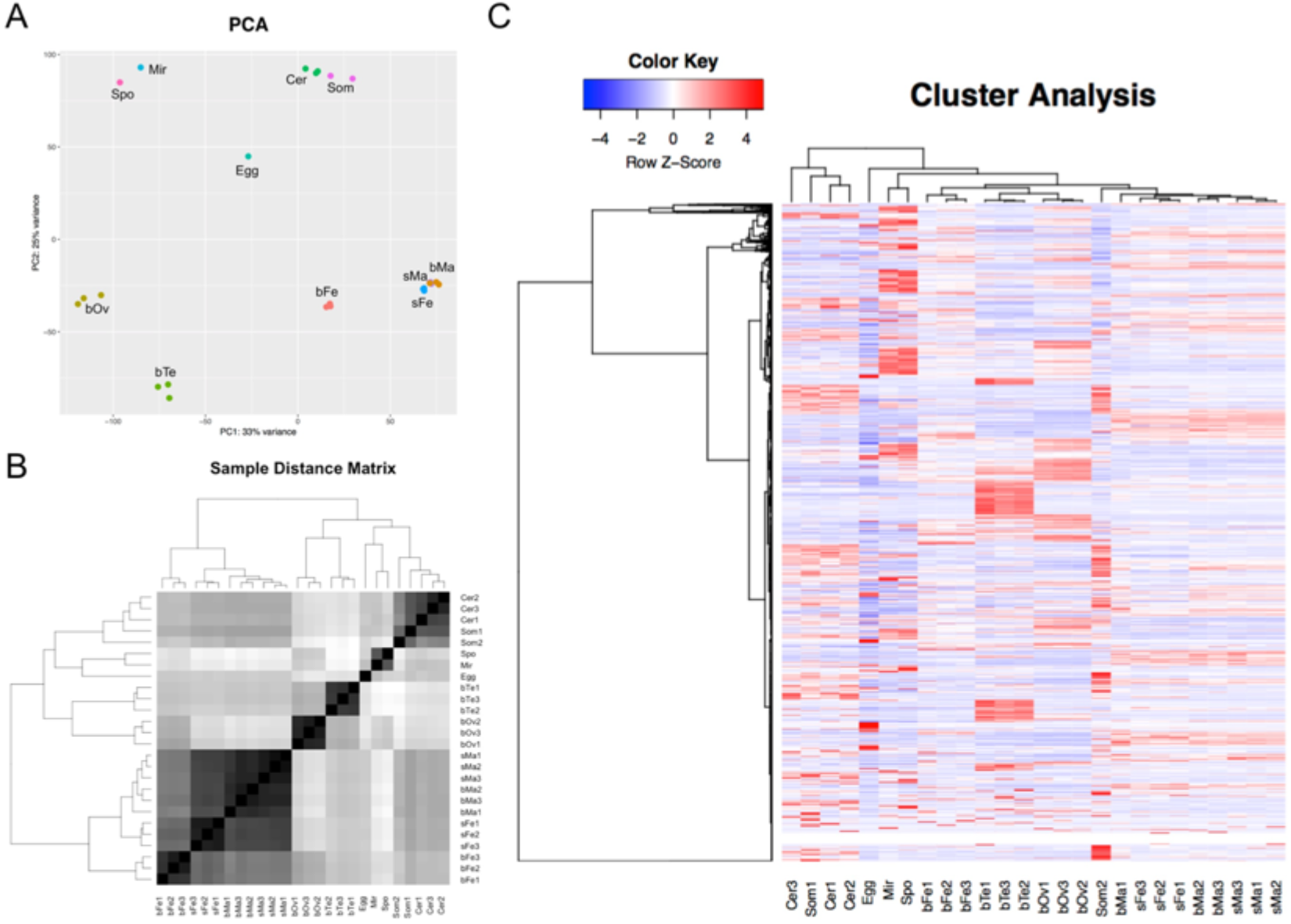
Visualisation of sample relationships using three different approaches. (A) Principle component analysis. Each dot represents a biological sample. (B) Sample distance matrix as revealed by a heat map. Sample clusters were indicated by the dendrogram. (C) Cluster analysis on both samples and genes. Note that although separated from Som1, Som2 showed similar expression patterns but at higher levels. bOv: ovary from paired female, bTe: testis from paired male, Mir: Miracidium, Spo: Sporocyst, Cer: Cercaria, Som: Schistosomulum, sFe: unpaired female, sMa: unpaired male, bMa: paired male, bFe: paired female.

**Table 2.** Summary of sample library size and transcriptome coverage

| Sample | Group | Library Size | No. genes with RPKM >1 | Coverage |
| --- | --- | --- | --- | --- |
| bTe1 | bTe | 29,643,786 | 7,904 | 78.21% |
| bTe2 | bTe | 24,625,567 | 7,895 | 78.12% |
| bTe3 | bTe | 19,939,503 | 7,940 | 78.57% |
| bOv1 | bOv | 24,831,879 | 7,284 | 72.08% |
| bOv2 | bOv | 63,472,787 | 7,135 | 70.60% |
| bOv3 | bOv | 56,664,560 | 7,115 | 70.40% |
| Egg | Egg | 96,691 | 5,855 | 57.94% |
| Mir | Mir | 60,230,741 | 7,010 | 69.36% |
| Spo | Spo | 66,122,608 | 6,737 | 66.66% |
| Cer1 | Cer | 14,990,523 | 7,735 | 76.54% |
| Cer2 | Cer | 32,609,995 | 7,977 | 78.93% |
| Cer3 | Cer | 23,453,370 | 7,932 | 78.49% |
| Som1 | Som | 20,887,726 | 8,173 | 80.87% |
| Som2 | Som | 27,539,735 | 8,240 | 81.54% |
| sMa1 | sMa | 42,949,245 | 8,264 | 81.77% |
| sMa2 | sMa | 42,841,035 | 8,245 | 81.59% |
| sMa3 | sMa | 36,581,319 | 8,312 | 82.25% |
| sFe1 | sFe | 43,913,097 | 8,337 | 82.50% |
| sFe2 | sFe | 75,421,480 | 8,340 | 82.53% |
| sFe3 | sFe | 50,058,027 | 8,318 | 82.31% |
| bMa1 | bMa | 52,977,773 | 8,403 | 83.15% |
| bMa2 | bMa | 40,595,996 | 8,381 | 82.93% |
| bMa3 | bMa | 40,319,774 | 8,306 | 82.19% |
| bFe1 | bFe | 15,839,979 | 8,154 | 80.68% |
| bFe2 | bFe | 29,547,879 | 7,945 | 78.62% |
| bFe3 | bFe | 25,180,870 | 7,943 | 78.60% |

**Based on the revealed sample clusters**, we obtained 407, 2,141, 1,955, 1,541, and 870 genes with preferential expression in Egg, Mir-Spo, Cer-Som, sM-sF-bM, and bFe, respectively (Supplementary Table 1). These preferentially-expressed genes make up 68% of *S. mansoni* protein-coding genes, and their product information indicates that they are responsible for the primary role of the parasite at specific life stage(s) (Fig. 2, Supplementary Table 1). For instance, splicing factors, RNA helicases, eIFs are commonly known as important for proliferation (Jankowsky 2011; Jackson et al. 2010), which happens tremendously in sporocyst; GPCRs, channel proteins and calcium-associated proteins are associated with sensory and motion (Lodish et al. 2000; Gover et al. 2009; Julius and Nathans 2012), important processes for cercaria and schistosomulum development; tetraspanins, tegument allergen, VALs and MEGs are well known to be involved in host-parasite interaction (Philippsen et al. 2015); and eggshell proteins are required for reproduction in *S. mansoni* (note that the number of eggshell proteins should be higher than indicated as many of them are annotated as hypothetical proteins). Top pathways associated with these genes were shown at the right on Fig. 2, which also supports the transition of dominant functions in the parasite.

**Figure 2.**
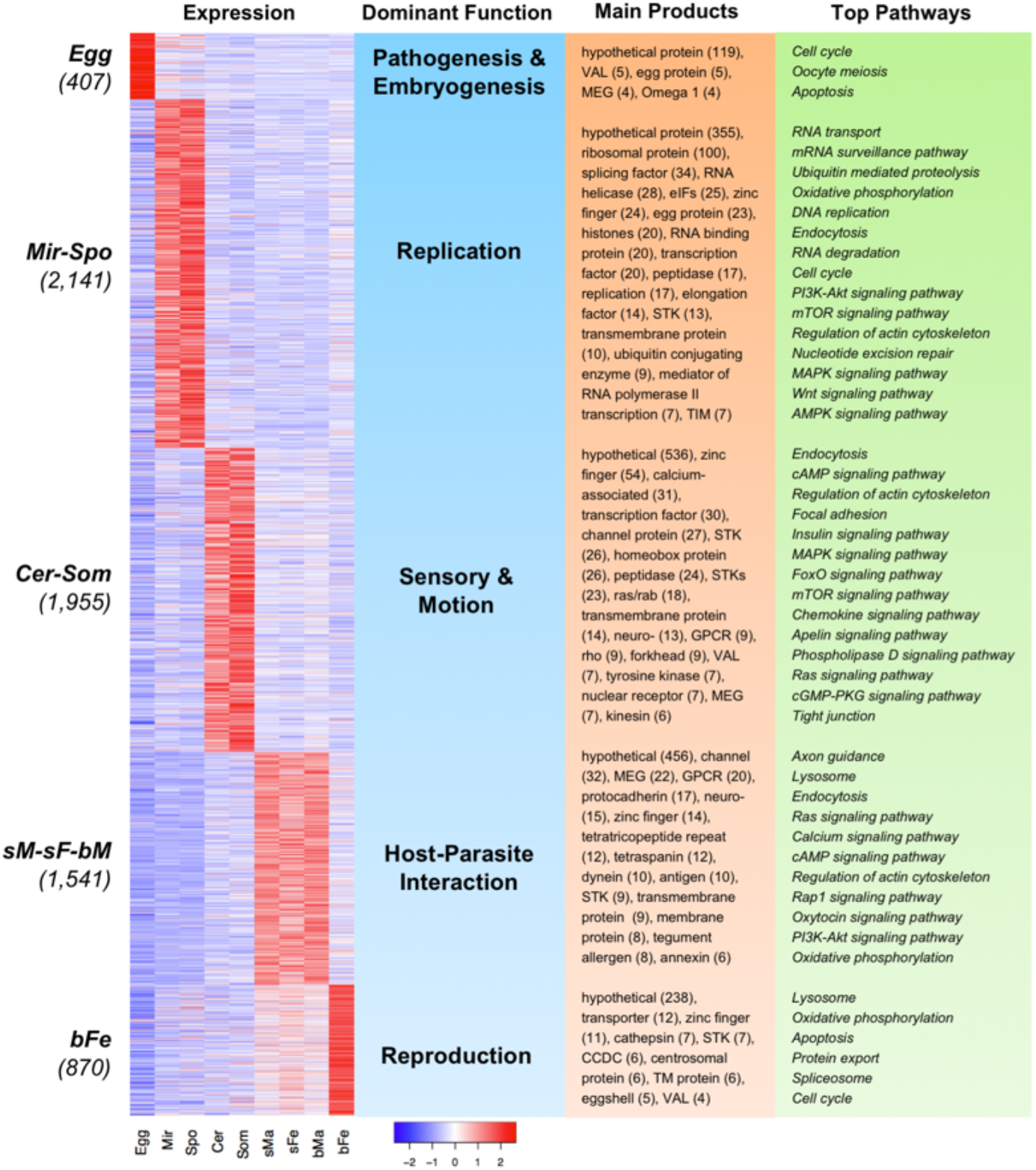
Preferentially-expressed genes and associated functions. Heat map of genes with preferential expression in certain life stages, the primary function of that stage, main products for these genes, and associated pathways were shown. Number of preferentially-expressed genes in each group was shown in the parentheses. Heat map was generated using the Z-score method (scaling on row). In the product information, number of genes coding for the same product was indicated in parentheses. For full list of products, please refer to Supplementary Table 1. In the pathway map, notations for common pathways, such as “Metabolic pathways” and “Ribosome”, or ambiguous pathways, such as “Huntington's disease” and “Vibrio cholerae infection”, were excluded from the list.

Top 10 genes from each of above enriched groups were summarised in Table 3 with information from previous investigations. The expression patterns of these genes are provided in Fig. 3A. Note that many of them are annotated as “hypothetical protein” (also see Fig. 2) and the patterns of their expression abundance will facilitate their predicted functions in the parasite, or improve the function annotation in the database (e.g., Smp_032670, Smp_193380 and Smp_179420 were annotated as “egg protein” but they have highest and preferential expression in mircidium / sporocyst).

**Figure 3.**
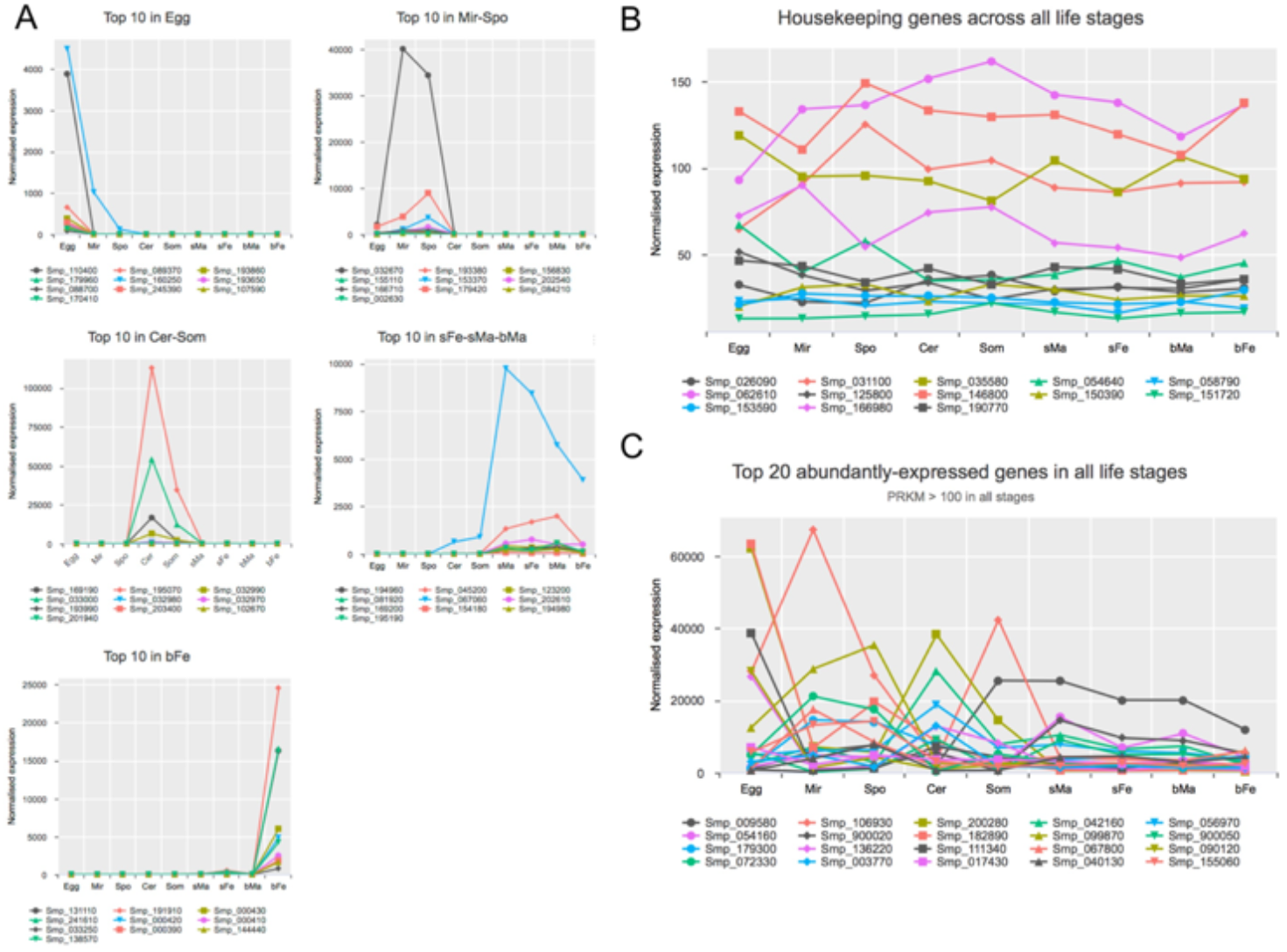
Expression profiles of exemplary genes. (A) Top 10 preferentially-expressed genes in certain life stage(s). (B) Housekeeping genes. (C) Top 20 abundantly-expressed genes in all life stages.

**Table 3.** Top 10 genes from each group

| *Preferentially-expressed in Egg* |  |  |  |
| --- | --- | --- | --- |
| Gene ID | Fold difference to the rest stages | Product description | Studies in life stage and reference |
| Smp_110400 | 303,218.9 | hypothetical protein | -- |
| Smp_089370 | 59,888.9 | Cell wall integrity and stress response | -- |
| Smp_193860 | 43,237.6 | hepatotoxic ribonuclease omega 1 | Egg (Dunne et al. 1991) |
| Smp_179960 | 21,618.8 | hepatotoxic ribonuclease omega 1 | -- |
| Smp_160250 | 18,951.2 | venom allergen (val) protein | NA (Chalmers et al. 2008) |
| Smp_193650 | 15,500.2 | mucin | -- |
| Smp_088700 | 12,416.8 | hypothetical protein | -- |
| Smp_245390 | 11,993.8 | IL-4-inducing protein | -- |
| Smp_107590 | 11,910.9 | hypothetical protein | -- |
| Smp_170410 | 9,607.9 | NADH:ubiquinone oxidoreductase complex I | Egg (Mathieson and Wilson 2010) |
| *Preferentially-expressed in Mir-Spo* |  |  |  |
| Gene ID | Fold difference to the rest stages | Product description | Studies in life stage and reference |
| Smp_032670 | 716,198.5 | egg protein C122 | Miracidium/sporocyst (Wu et al. 2009) |
| Smp_193380 | 55,492.3 | egg protein CP391S | -- |
| Smp_156830 | 40,905.3 | hypothetical protein | -- |
| Smp_155110 | 26,615.9 | voltage gated potassium channel | -- |
| Smp_153370 | 19,484.0 | hypothetical protein | -- |
| Smp_202540 | 19,349.4 | hypothetical protein | -- |
| Smp_166710 | 17,559.9 | hypothetical protein | -- |
| Smp_179420 | 10,085.5 | egg protein CP391S | -- |
| Smp_084210 | 9,345.1 | gag pol polyprotein | -- |
| Smp_002630 | 7,383.0 | venom allergen-like (VAL) 2 protein | Miracidium/Sporocyst (Chalmers et al. 2008; Wu et al. 2009) |
| *Preferentially-expressed in Cer-Som* |  |  |  |
| Gene ID | Fold difference to the rest stages | Product description | Studies in life stage and reference |
| Smp_169190 | 513,498.3 | tegument-allergen-like protein | Cercaria/Schistosomula (Fitzsimmons et al. 2012; Gava et al. 2017);<br>Adult male (Leutner et al. 2013) |
| Smp_195070 | 31,216.0 | cercarial stage specific protein Sj20H8 | -- |
| Smp_032990 | 25,531.7 | Calmodulin 4 (Calcium binding protein Dd112) | Adult female (Buro et al. 2013; Lu et al. 2015);<br>Schistosomulum (Nowacki et al. 2015) |

| **Smp_033000** | 10,960.3 | calcium-binding protein | Adult male (Leutner et al. 2013) |
| --- | --- | --- | --- |
| **Smp_032980** | 6,038.6 | calmodulin protein | Schistosomulum (Nowacki et al. 2015) |
| **Smp_032970** | 2,486.7 | calmodulin protein | -- |
| **Smp_193990** | 996.0 | hypothetical protein | -- |
| **Smp_203400** | 929.3 | rhodopsin orphan GPCR | -- |
| **Smp_102670** | 916.5 | hypothetical protein | -- |
| **Smp_201940** | 916.5 | hypothetical protein | -- |
| ***Preferentially-expressed in sM-sF-bM*** |  |  |  |
| **Gene ID** | **Fold difference to the rest stages** | **Product description** | **Studies in life stage and reference** |
| **Smp_194960** | 8,964.5 | 25 kDa integral membrane protein | -- |
| **Smp_045200** | 4,329.5 | tegument-allergen-like protein | Schistosomulum and adult tegument (Fitzsimmons et al. 2012) |
| **Smp_123200** | 4,124.5 | MEG-32.2 protein | Adult head (Wilson et al. 2015) |
| **Smp_081920** | 3,396.9 | hypothetical protein / CD59-like | Adult tegument (Collins et al. 2016) |
| **Smp_067060** | 2,574.4 | Cathepsin B1 isotype 2 | Adult vomitus (Philippsen et al. 2015) |
| **Smp_202610** | 1,820.4 | hypothetical protein | -- |
| **Smp_169200** | 1,584.7 | tegument-allergen-like protein | Adult (Fitzsimmons et al. 2012) |
| **Smp_154180** | 1,584.7 | 25 kDa integral membrane protein | Orthologue in *S. japonicum* tegument (Wu et al. 2011) |
| **Smp_194980** | 1,478.6 | 25 kDa integral membrane protein | Schistosomulum (Parker-Manuel et al. 2011) |
| **Smp_195190** | 1,305.2 | 13 kDa tegumental antigen Sm13 | Adult tegument (Collins et al. 2016; Wilson 2012) |
| ***Preferentially-expressed in bFe*** |  |  |  |
| **Gene ID** | **Fold difference to the rest stages** | **Product description** | **Studies in life stage and reference** |
| **Smp_131110** | 122,294.5 | hypothetical protein (Sm_p14) | Female vitellarium (Bobek et al. 1988) |
| **Smp_191910** | 49,667.0 | Stress protein DDR48 | Male oesophagus (Nawaratna et al. 2014); Female (Fitzpatrick et al. 2009) |
| **Smp_000430** | 29,125.6 | eggshell protein | Female vitellarium (Buro et al. 2013) |
| **Smp_241610** | 14,562.8 | P48 eggshell protein | Female vitellarium (Chen et al. 1992) |
| **Smp_000420** | 12,677.7 | Pro His rich protein | Male oesophagus (Nawaratna et al. 2014) |
| **Smp_000410** | 12,590.1 | Trematode Eggshell Synthesis | Male oesophagus (Nawaratna et al. 2014) |
| **Smp_033250** | 12,245.8 | hypothetical protein | -- |
| **Smp_000390** | 12,161.2 | hypothetical protein | Female (Anderson et al. 2015) |
| **Smp_144440** | 11,113.3 | replication A protein | -- |
| **Smp_138570** | 10,297.5 | spore germination protein | -- |

### Distinct differential expression patterns occur during parasite development

To obtain detailed gene expression during parasite development, and to validate the results of meta-analysis, pairwise differential gene expression analysis was performed for adjacent life stages. By setting the threshold at FDR < 0.01 and fold difference > 2, the number of differentially expressed genes (DEGs) were summarised in Fig. 4 (See detailed lists in Supplementary Table 2). We can observe a massive change in gene expression levels from sporocyst to cercariae, and from schistosomulum (skin-stage) to adult (numbers of DEGs > 4,300), whereas less DEGs were obtained in other comparisons. The results obtained by the meta-analysis were in good correlations with previous individual studies in either the larval stages (e.g., Spo vs Mir (Wang et al. 2013), Som vs Cer (Protasio et al. 2012)), or in the adult stage (e.g., bTe vs bOv, and adult comparisons between male and female or before and after pairing (Lu et al. 2016)) (Fig. 5).

**Figure 4.**
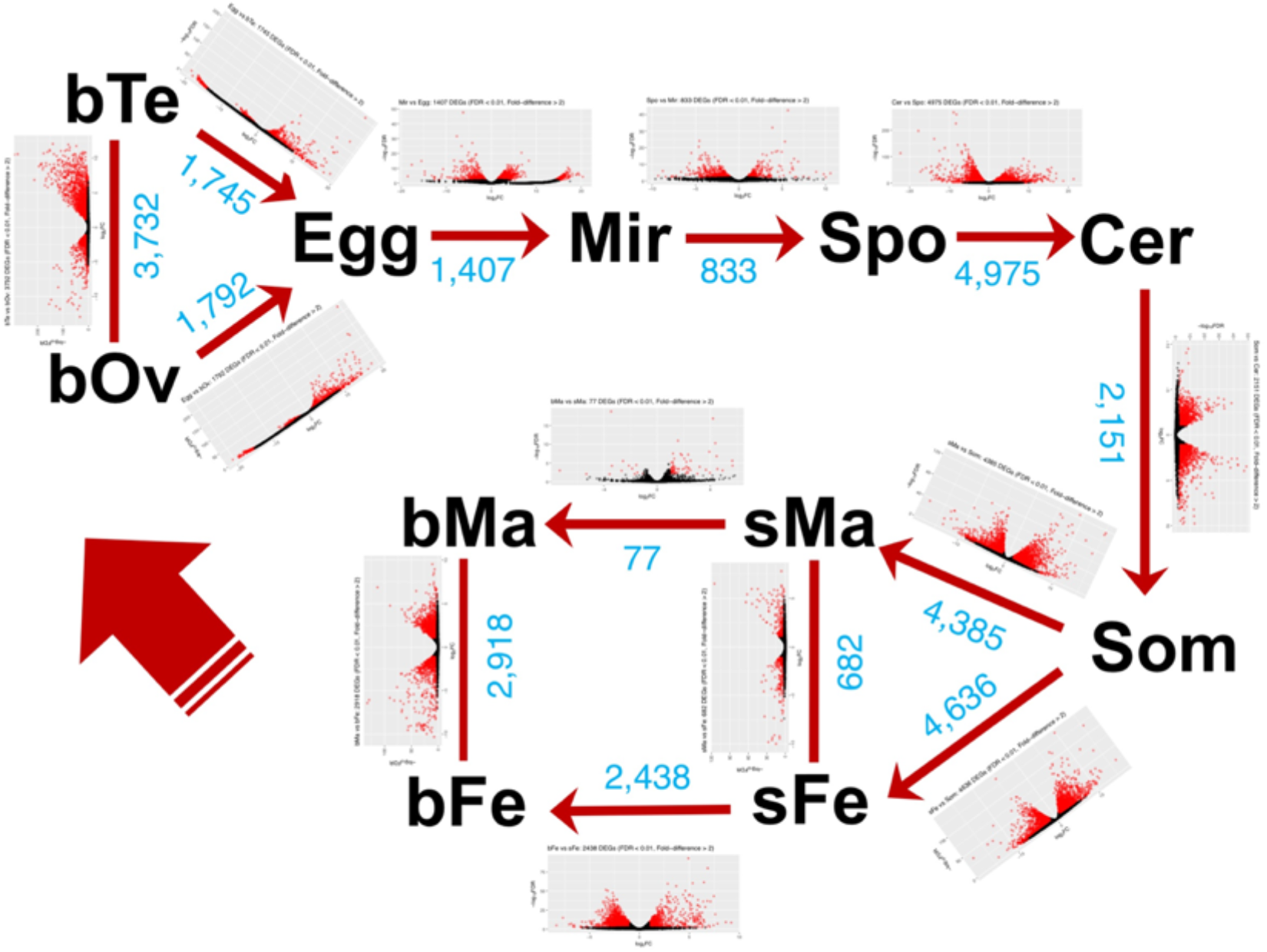
Numbers of differentially expressed genes between adjacent life stages. Total numbers of DEGs and log_2_FC-log_10_FDR plot with DEGs in red were included. Threshold: FDR < 0.01; fold-difference > 2.

**Figure 5.**
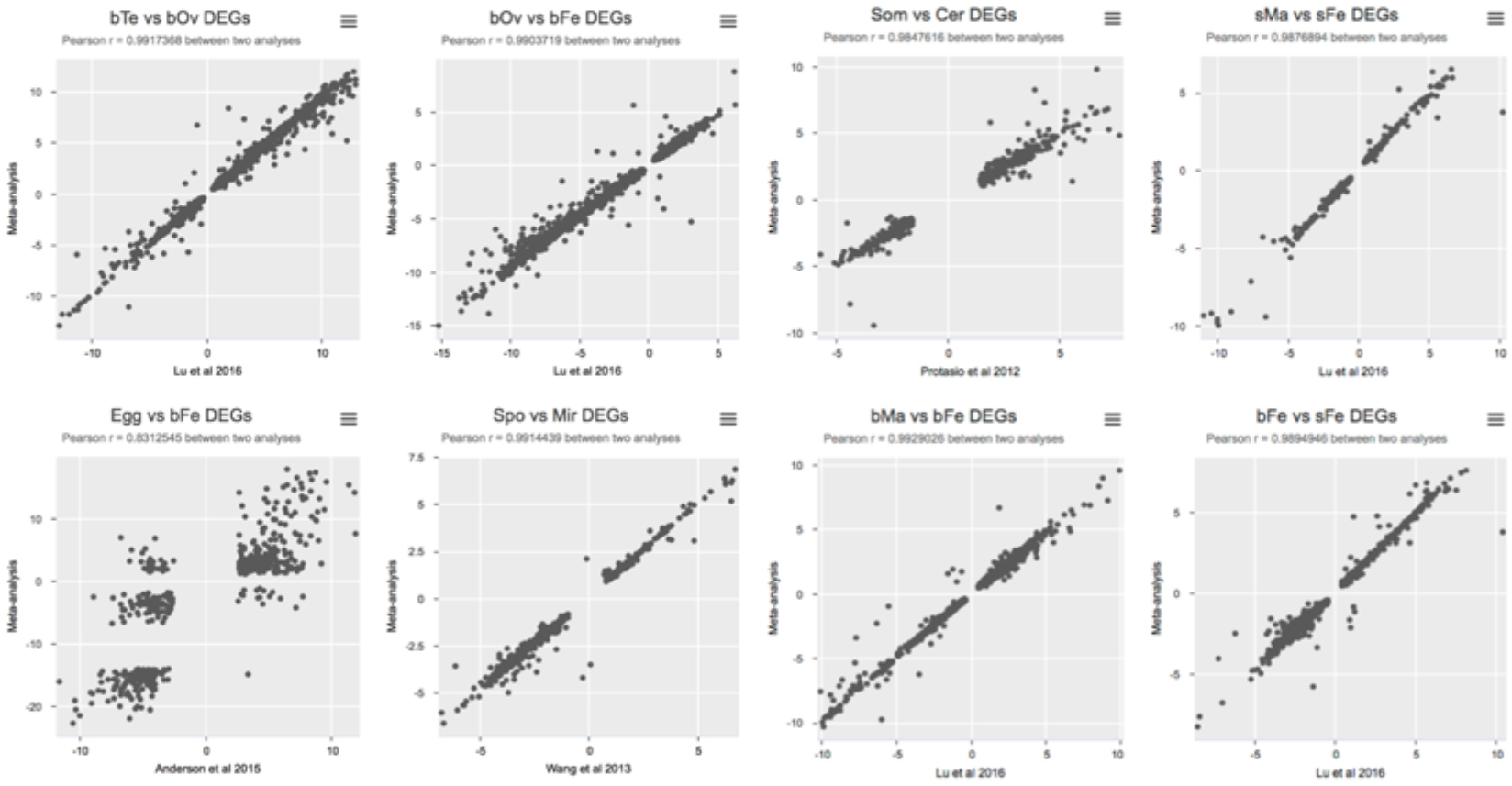
Log_2_FC correlations between meta-analysis and original analyses. Pearson’s correlation coefficient was indicated in each plot.

Besides the known comparisons, we can also obtain new information about the transitions from gonads to the egg/embryo, from sporocyst to cercaria, as well as from schistosomula to adults (Supplementary Table 2). The transitions not only reflect in the morphological change, but also in changing the biological processes, as reflected in significantly enriched GO terms in each stage. The mostly significantly enriched GO terms in sporocyst include mRNA splicing, RNA secondary structure unwinding, mitotic cell cycle process, and regulation of translation, whereas in cercaria they include cilium assembly, transport, chemical synaptic transmission, cell surface receptor signalling pathway (Fig. 6A). This observation is consistent with our previous conclusions (Fig. 2). As for the transformation from skin-stage schistosomulum to adult worm, the GO terms also reveal different processes. In schistosomulum, abundantly-expressed genes are involved in positive regulation of transcription, cell differentiation, anatomical structure development, response to lipid, etc, which indicate early response to host factors. After developed into adult worm, the processes switch to cilium assembly, oxidation-reduction process, transport along microtubule, ATP metabolic process, etc (Fig. 6B).

**Figure 6.**
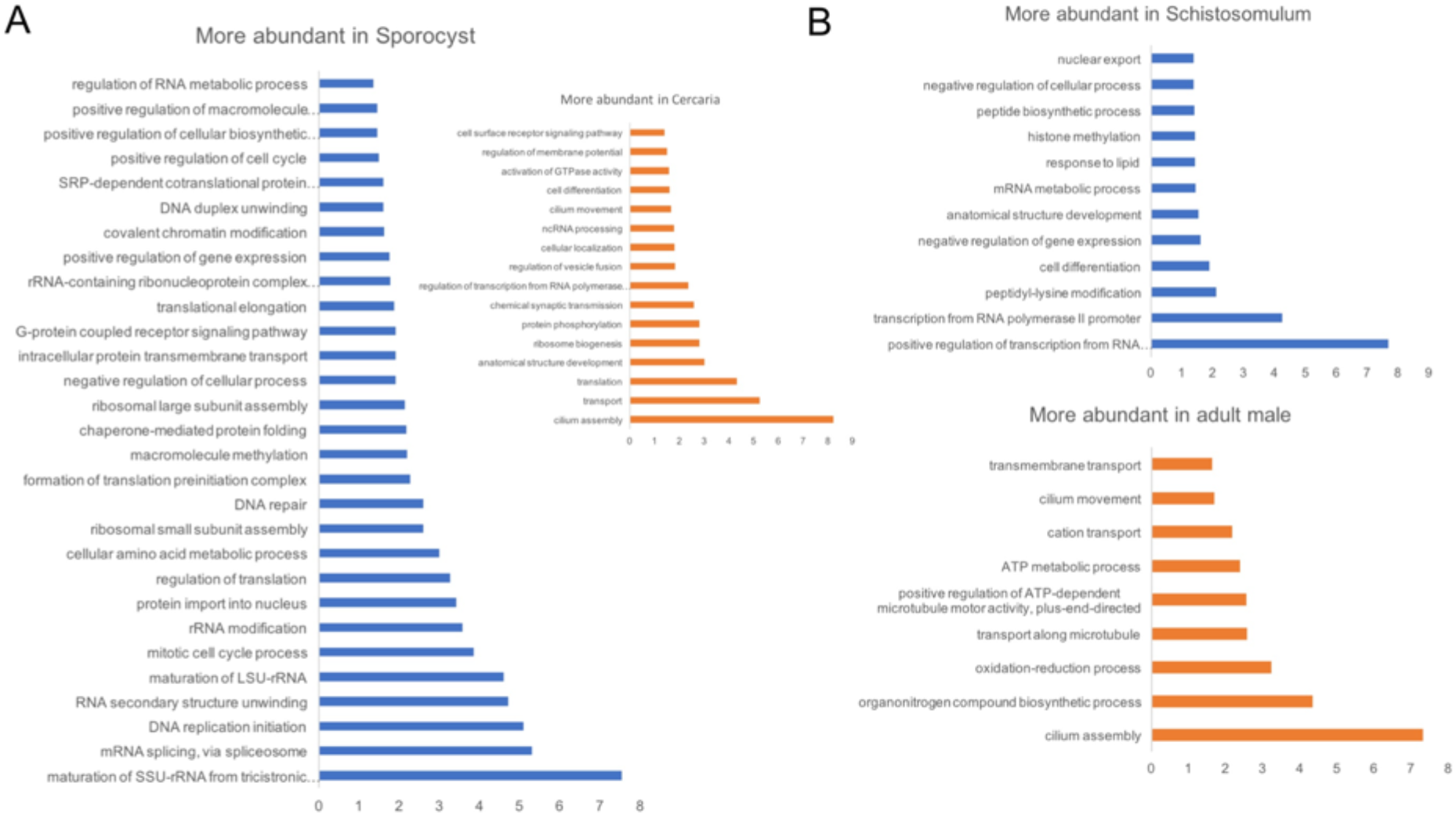
GO enrichment for differentially expressed genes. (A) DEGs between sporocyst and cercaria. (B) DEGs between schistosomulum and adult male. P < 0.05 for significant enrichment. Axis values represent -log_10_(P-Value) values.

### Novel housekeeping genes were identified across all life stages

Thirteen potential housekeeping genes were identified by the GLM approach (excluding gonads). Furthermore, their suitability was tested by calculating the stability value among all samples using Normfinder (Andersen et al. 2004). The gene IDs, product information, and stability values were summarised in Table 4. Expression patterns can be seen in Fig. 3B.

Most of them demonstrate good stabilities, and are suitable candidates as ubiquitously- and/or strongly-expressed genes.

**Table 4.** Housekeeping genes across all life stages

| Gene ID | Product | Stability |
| --- | --- | --- |
| **Smp_146800** | atlastin 2 | 0.12 |
| **Smp_035580** | serine:threonine protein phosphatase PP1 beta | 0.16 |
| **Smp_031100** | liquid facets | 0.17 |
| **Smp_058790** | mitogen activated protein kinase | 0.17 |
| **Smp_062610** | autophagy protein 101 | 0.17 |
| **Smp_026090** | ras gtp binding protein d | 0.18 |
| **Smp_153590** | tubulin specific chaperone D | 0.19 |
| **Smp_150390** | WD repeat containing protein 48 | 0.20 |
| **Smp_125800** | 26S proteasome non ATPase regulatory subunit | 0.22 |
| **Smp_190770** | lanC protein 2 like | 0.22 |
| **Smp_151720** | ubiquitin associated and SH3 domain containing protein | 0.24 |
| **Smp_054640** | major facilitator superfamily domain containing protein | 0.25 |
| **Smp_166980** | transcription initiation factor TFIID subunit | 0.25 |

### Most abundantly-expressed genes in all life stages

Abundantly-expressed genes can be used for functional genomics studies, e.g., promoters for transgenes, or for screening targets for vaccine development. With ranked average expression in all stages (excluding gonads) and to avoid zero-inflation, we select those with RPKM > 100 in all samples as ubiquitously abundant genes. Table 5 shows a summary for top 20 of those genes, and their expression profiles can be obtained from Fig. 3C. Some of them are previously known as ubiquitously-expressed genes, e.g., HPS70 (Smp_106930) (Neumann et al. 1993), and as vaccine candidates, e.g., aldolase (Smp_042160), GAPDH (Smp_056970), GTS28 (Smp_054160) (Wilson et al. 2016).

**Table 5.** Top 20 abundantly-expressed genes in all life stages

| Gene ID | Product | Avg expr (rpkm) |
| --- | --- | --- |
| **Smp_009580** | polyubiquitin ubiquitin | 15,816.32 |
| **Smp_106930** | heat shock 70 kDa protein homolog | 10,617.61 |
| **Smp_200280** | hypothetical protein | 9,910.12 |
| **Smp_042160** | fructose-bisphosphate aldolase | 8,750.15 |
| **Smp_056970** | glyceraldehyde 3 phosphate dehydrogenase | 7,113.44 |
| **Smp_054160** | glutathione S-transferase class-mu 28 kDa isozyme | 6,923.32 |
| **Smp_900020** | NADH dehydrogenase subunit 6 | 6,862.36 |
| **Smp_182890** | hypothetical protein | 5,517.16 |
| **Smp_099870** | elongation factor 1-alpha | 5,502.19 |
| **Smp_900050** | NADH dehydrogenase subunit 5 | 4,982.15 |
| **Smp_179300** | cellular nucleic acid binding protein | 4,568.08 |
| **Smp_136220** | hypothetical protein | 4,501.24 |
| **Smp_111340** | hypothetical protein | 4,498.75 |
| **Smp_067800** | hypothetical protein | 4,025.97 |
| **Smp_090120** | alpha tubulin | 3,278.92 |
| **Smp_072330** | heat shock protein heat shock protein 86 | 3,028.51 |
| **Smp_003770** | histone H1 | 3,010.62 |
| **Smp_017430** | multivalent antigen sj gapdh | 3,005.27 |
| **Smp_040130** | cyclophilin peptidyl-prolyl cis-trans isomerase | 2,975.11 |
| **Smp_155060** | phosphatase 2a inhibitor i2pp2a | 2,960.76 |

## Discussion

Meta-analysis of gene expression has been exploited in other organisms such as humans, either with RNA-seq data across species and tissues (Sudmant et al. 2015), or with Microarray data from different studies (O’Mara et al. 2016). While technical differences and additional biological variabilities might exist among studies, a robust normalisation and differential analysis method is critical for reliable results. Trimmed Mean of M-values (TMM) is a batch normalisation method on a group of samples and has been proven to perform well in bulk RNA-seq (Dillies et al. 2013). As for differential expression analysis, the negative binomial generalized linear model (GLM) was tested suitable for low inter-study variation and small numbers of studies (Rau et al. 2014). In our case, by applying the TMM normalisation and GLM approach for differential expression analysis, we obtained reliable results having very good correlations with original studies. In the case of the egg sample, the variability was probably due to technical difference (*de novo* assembly vs reference mapping) and natural differences between *S. mansoni* strains (Brazilian vs Liberian strain).

Genes identified to be preferentially expressed in certain life stages can be valuable resource for the research community. On one hand, it can be a guidance for investigating gene functions in proper life stage(s), as many genes show extremely higher expression in one stage than in the others, e.g. calmodulins in cercariae. On the other hand, stage-preferential gene expression can benefit functional gene annotations. For instance, there are 23 genes annotated as “egg protein” but detected in our analysis as preferentially-expressed in miracidium/sprocyst. Another example is that many hypothetical proteins show preferential expression in bFe (paired female; Fig. 2), and by comparing the expression in whole female with that in the ovary, we can probably estimate the gene function in the vitellarium.

Finally, many of the identified genes were consistent with previous studies (Table 3), indicating the robustness of the meta-analysis.

Our analysis also confirms assumptions from previous studies. For example, *elav2* (Smp_194950) and *cdc25* (Smp_152200) were previously found to be exclusively transcribed in testes in the adult stage (Lu et al. 2016), and our analysis extended the data and supported the conclusion (Fig. 7A). Furthermore, *cpeb1* (Smp_070360) was proposed to fulfil roles in oocyte maturation (Lu et al. 2016; Wang et al. 2017), and the meta-analysis suggests its additional function in the embryo (Fig. 7A), as a dual-function protein also found in *Xenopus* (Novoa et al. 2010). In addition, our data supports the differential expression of aromatic-L-amino-acid decarboxylase (AADC) and allatostatin-A receptor-like gene (AlstR) between male and female worms, which were characterised in a recent study in *Schistosoma japonicum* as mediator of reproduction (Wang et al. 2017). Their *S. mansoni* orthologues exhibiting similar expression patterns (Fig. 7B) indicates a similar regulation theme.

**Figure 7.**
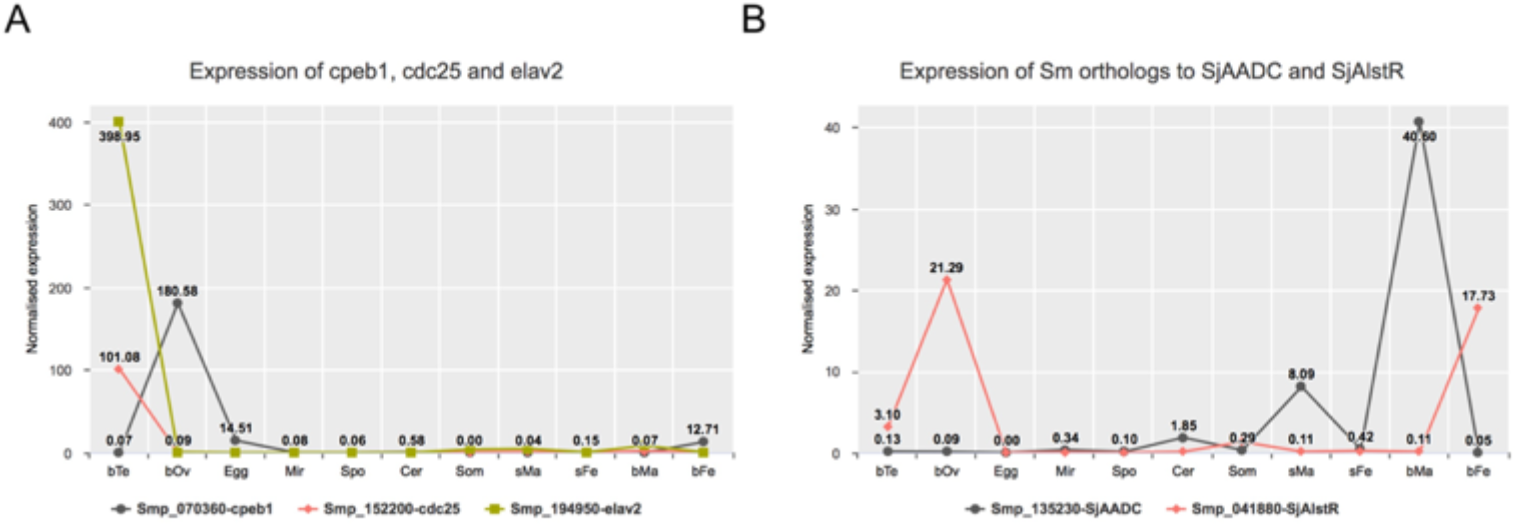
Expression profiles of selected genes highlighted in previous studies. (A) Elav2, cdc25, and cpeb1 show preferential expression in gonads; (B) *S. mansoni* orthologues of *SjAADC* and *SjAlstR* exhibit similar differential expression patterns in male and female worms as *in S. japonicum*.

When we calculated differential gene expression between adjacent stages, we obtained quite distinct numbers of DEGs (Fig. 4). The most significant gene expression changes were observed during the transformation from sporocyst to cercaria, as well as from schistosomulum (3h skin stage) to adult worm. The former can be associated with the changes from a less active state where the parasites are mainly manipulating themselves, to a state where they need to swim fast, and to sense the definitive host efficiently, as confirmed in GO enrichment analysis (Fig. 6A). We saw that genes more abundantly-expressed in sporocyst are mainly involved in cell replication (e.g., mRNA splicing, DNA replication initiation, translational elongation, etc.), and those in cercaria are mainly involved in sensory and mobility (e.g., cilium assembly / movement, transport, chemical synaptic transmission, etc.) The latter can reveal many physical and biochemical changes in the parasite from skin- to lung-stage, accompanied by increased immunological defences (GOBERT et al. 2007). Overall, the tremendous differential expression is associated with the morphological and functional changes in theses stages, and can support our understanding of schistosome developmental biology.

With respect to venom allergen-like (VALs) genes, our data supports previous findings. As show in Fig. 8A, SmVALs are distributed in different life stages. While consistent with previous finding on the presence of SmVALs2, 3, 5, 9 in egg, miracidia and sporocyst (Chalmers et al. 2008), we found the highest abundance in the egg instead of miracidia stage as discovered by the authors. This is probably due to the unsuitability of the housekeeping gene used in the previous study. The same expression patterns were found for SmVAL23 and SmVAL29, as discovered before (Wu et al. 2009). In addition, we supported the discovery of SmVAL22 preferential expression in sporocyst (Wang et al. 2013). SmVAL1 and SmVAL24 were found to be nearly exclusively in cercaria, which differs from the finding about higher SmVAL24 expression in the germ balls (Fernandes et al. 2017). More interestingly, we discovered two VALs that seem exclusive for the testis, whose functions need further investigations.

**Figure 8.**
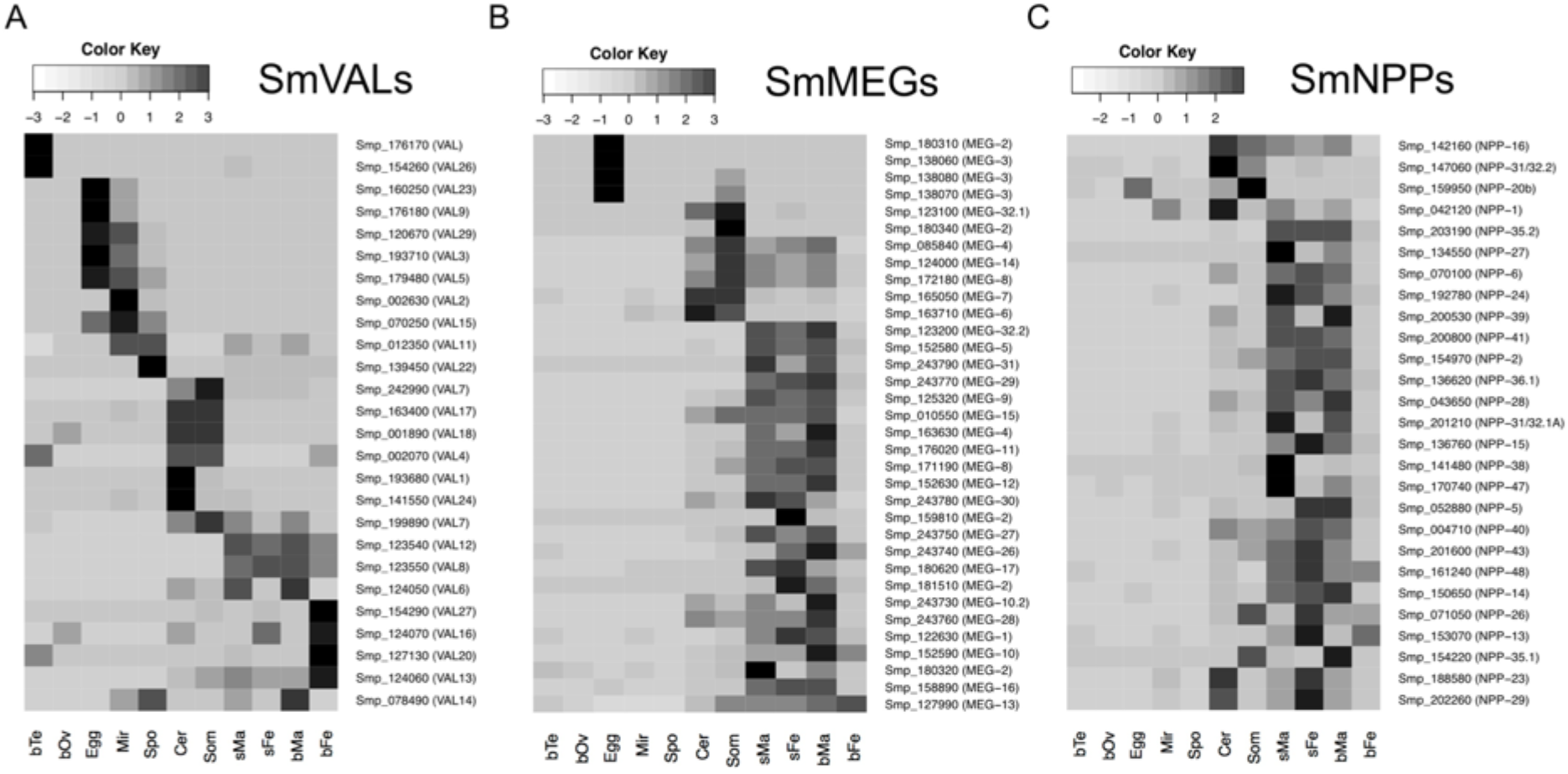
Heat maps reveal relative expression abundance in all samples. (A) SmVALs. (B) SmMEGs. (C) SmNPPs. VAL: venom allergen-like; MEG: micro exon gene; NPP: neuropeptide precursor.

Micro-exon genes (MEGs) were previously reported to be clustered in the esophagus of adult schistosomes (Wilson et al. 2015; Wang and Collins 2016) and proposed with functions for blood processing. Our data supports that by showing that most of known SmMEGs were found to be preferential in the sM-sF-bM group (Fig. 8B). We also identified that some members of SmMEG2 and SmMEG3 are preferential in the egg, which agrees with previous comparison between expression of them in egg to cercaria (DeMarco et al. 2010).

Our results also give new information about expression of neuropeptides in schistosomes. Based on the discovery of novel neuropeptide precursors (NPPs) in flatworms (Koziol et al. 2016), we found that most of SmNPPs were preferentially-expressed in sM-sF-bM (Fig. 8C), a pattern identified recently in the adult stage (Lu et al. 2016). This indicates active neuronal processes at the host-parasite interface.

The identified novel housekeeping genes and abundantly-expressed genes can be useful for quantitative and/or functional studies. While candidates for housekeeping genes had been proposed before (Lu et al. 2016), the presented comprehensive analysis can provide more reliable result as it covers all life stages of schistosomes. From the expression data they seem to have a reasonable transcript abundance and can be used for testing purposes (Fig. 3B). Abundantly-expressed genes normally exhibited strong promoter activities thus suitable for functional approaches including overexpression or know-down of specific genes, and for vaccine development. We extracted 20 strongly expressed genes in all life stages (Table 5; Fig. 3C). While some of them have been discussed before, the list provides new candidates for such purposes.

## Data Availability

All RNA-seq data analysed in this work can be obtained from the European Nucleotide Archive (http://www.ebi.ac.uk/ena). Accession numbers were summarised in Table 1. Tools and parameters used in the analysis can be found in the methods section. Gene expression profile can be accessed via https://meta.schisto.xyz

## Author contributions

Zhigang Lu

Roles: Conceptualization, Data Curation, Formal Analysis, Resources, Methodology, Visualisation, Writing - Original Draft Preparation

Matthew Berriman

Roles: Formal Analysis, Writing - Review & Editing

## Supplementary Material

Supplementary Table 1. List of genes with abundant expression in different life stages and their product information (.csv file).

Supplementary Table 2. Log_2_FC and FDR values originated from various differential expression analyses (.csv file).

